# Estimating numbers of fluorescent molecules in single cells by analysing fluctuations in photobleaching

**DOI:** 10.1101/272310

**Authors:** Elco Bakker, Peter S. Swain

## Abstract

The impact of fluorescence microscopy has been limited by the difficulties of expressing measurements of fluorescent proteins in numbers of molecules. Absolute numbers enable the integration of results from different laboratories, empower mathematical modelling, and are the bedrock for a quantitative, predictive biology. Here we develop a general algorithm to infer numbers of molecules from fluctuations in the photobleaching of proteins tagged with Green Fluorescent Protein. To untangle measurement noise from stochastic fluctuations, we use the linear noise approximation and Kalman filtering within a framework of Bayesian inference. Not only do our results agree with biochemical measurements for multiple proteins in budding yeast, but we also provide a statistically verified model of measurement noise for fluorescence microscopes. The experiments we require are straightforward and use only a wide-field fluorescence microscope. As such, our approach has the potential to become standard for those practising quantitative fluorescence microscopy.

## Introduction

In fluorescence microscopy, converting measurements of fluorescence into numbers of molecules is a long-standing challenge. This deficit limits both the ability to combine fluorescence measurements from different experiments into one data set and the application of quantitative analyses of time-series that assumes the numbers of molecules are known [1, 2, 3, 4, 5].

Although methods exist to estimate the ratio between protein number and fluorescence, most require additional expertise and equipment beyond that typically needed for fluorescence microscopy [6]. Examples include fluorescence correlation spectroscopy [7, 8], image correlation spectroscopy [9], photon-counting histogram analysis [10], and fluorescence intensity distribution analysis [11], but all these approaches require at least a confocal microscope. Data is publicly available for alternative biochemical methods, such as quantitative Western blotting [12, 13] and mass spectrometry [14], but their accuracy is difficult to assess [15].

Ideally we would like to have a technique that requires only a wide-field fluorescence microscope and can be applied directly to the cells whose fluorescence is of interest. One promising set of approaches are fluctuation-based methods. These methods were first proposed in 1974 [7], but since then no standard approach has been developed for wide-field microscopy. Fluctuation analyses exploit that the magnitude of fluctuations in fluorescence are determined not by the concentration of the fluorescent molecules but by their numbers [16] and infer the fluorescence per molecule by analysing deviations from the mean (Fig. 1A).

**Figure 1.**
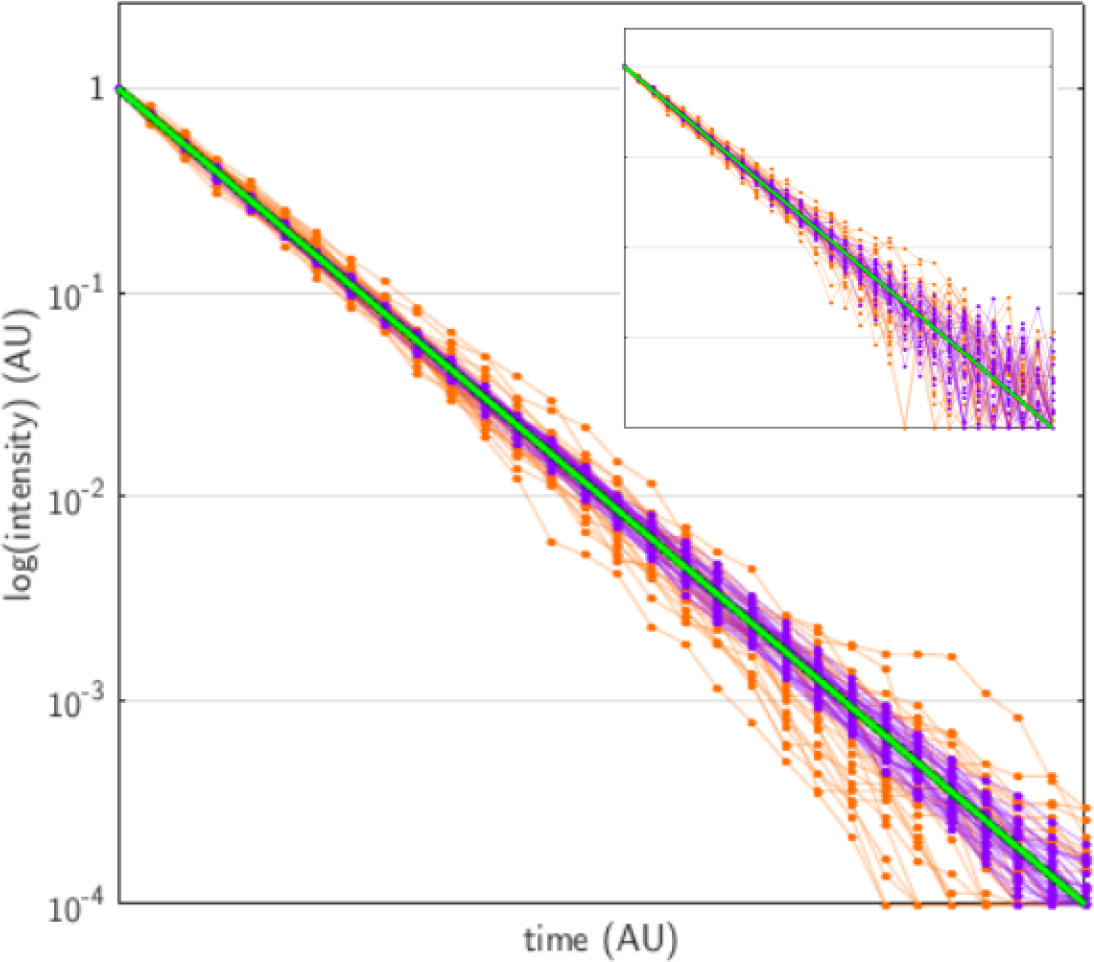
The magnitude of fluctuations is determined by numbers of molecules. Simulated time-series for stochastic exponential decay are normalized to the same starting value and plotted on a log scale for both low and high numbers of proteins. The mean behaviour is common (green line), but the data for low numbers of proteins (orange) shows larger deviations from the mean than the data for high numbers of proteins (purple). Inset: Stochastic fluctuations become masked with measurement noise (here an independent, additive Gaussian noise with zero mean and a standard deviation of 0.01): the orange and purple time-series now substantially overlap.

Bespoke fluctuation techniques have been developed for wide-field microscopes, which do not focus on diffusion as fluorescence correlation spectroscopy does but instead use other stochastic events. There are two main approaches. One is to study fluctuations in the distribution of fluorescent proteins between daughter cells at cell division [17, 18, 19]. These techniques have been mostly applied to bacteria, are unsuitable for non-dividing cells [20] and do not straightforwardly extend to species that exhibit differences in size between mothers and daughters. Further, the data must be obtained over several tens of divisions, which may be prohibitive for slowly dividing cells. The other approach is to study fluctuations in stochastic decay processes. Interventions to inhibit translation and transcription have allowed the fluorescence per molecule to be estimated in mammalian cells [4], but the stability of fluorescent proteins can make these experiments time consuming. A faster approach is to deliberately induce photobleaching [21]: the process whereby fluorophores cease to fluoresce when continuously excited [22]. This method has been applied in vitro [23, 24] and to bacteria [25], but the proposed analysis relies on photobleaching exhibiting an exponential decay [21], which is expected for single molecules but not for the fluorescence of cells [22].

Analysing such data is challenging. We must fit not the mean behaviour but fluctuations around the mean, and these fluctuations are typically confounded by measurement noise. To be successful, any inference procedure must be able to distinguish between measurement noise and stochastic effects (Fig. 1 inset).

Here we calibrate data from fluorescence microscopy through a Bayesian analysis of the fluctuations during photobleaching of eukaryotic cells. We model photobleaching as a multiexponential process [22] and include measurement noise, unlike earlier analyses [21, 23, 25, 24], which neglect both. Using budding yeast, we show that photobleaching is indeed not described by a single exponential decay and introduce, fit, and statistically compare models that differ in their description of measurement noise. We therefore find both an empirically verified distribution for the measurement noise present in wide-field fluorescence microscopy, which can be used in the analysis of other data, and the fluorescence per molecule for proteins tagged with Green Fluorescent Protein (GFP). The corresponding predictions of protein numbers agree with estimates found by biochemical methods for the six proteins tested.

## Results

### A biophysical model of photobleaching

We obtained time-series of photobleaching using budding yeast and GFP-tagged proteins [26]. Cells were fixed and photobleached in sustained illumination with fluorescence measurements taken every 10 seconds (Fig. 2 and Methods).

**Figure 2.**
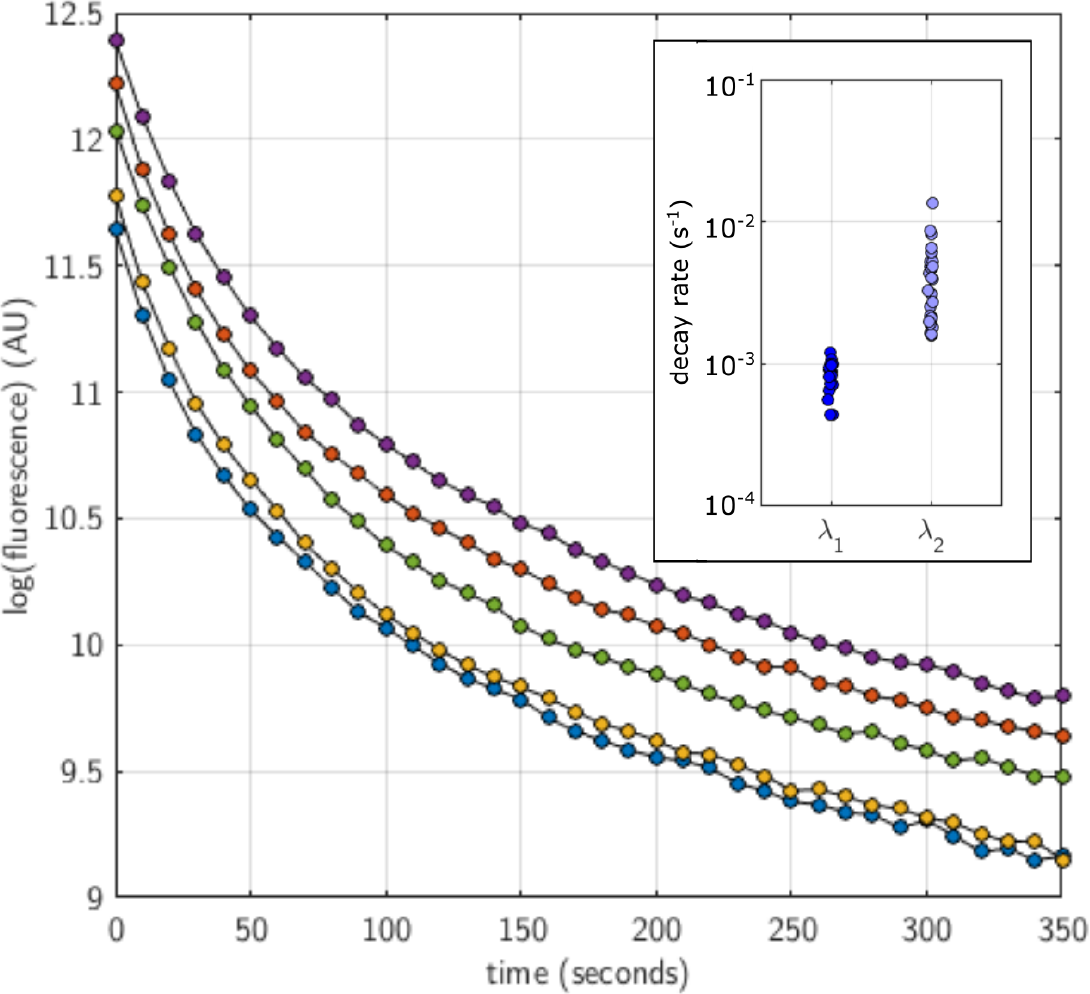
Photobleaching is not described by a single exponential decay. The logarithm of the fluorescence of 5 cells of budding yeast expressing Pgk1-GFP and undergoing photobleaching does not fall on a straight line as a function of time (in contrast to the data of Fig. 1). The inset shows all the pairs of decay rates found from fitting a bi-exponential decay to the time-series of each cell (using maximum likelihood). There is substantial cell-to-cell variation (each circle represents a cell and notice the log scale on the *y*-axis).

Our data is incompatible with earlier methodologies (Methods) because it is not well described by a single exponential decay (Fig. 2). First, the data for any individual cell shows systematic deviations from a single exponential and is better fit by a bi-exponential decay. Second, the decay rates of these two exponentials vary substantially between cells (Fig. 2 inset). Multi-exponential photobleaching is common [22, 27, 28, 29, 30] and can be caused by heterogeneous illumination [31], differing intracellular micro-environments [32], molecular rotation [33], and higher order interactions between excited fluorophores [34, 35]. These phenomena can also cause heterogeneity in parameters between cells if there is variation in either illumination or chemical composition across the population of cells.

We therefore consider each cell to have two populations of fluorescent proteins that photobleach with different rates. Writing the number of fluorescent proteins in cell *j* in the first population as **x**_1,*j*_ and the number of fluorescent proteins in the second population as x_2,*j*_, we model photobleaching as 
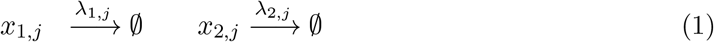
where λ_1,*j*_ is the rate of photobleaching of the first population in cell *j* and λ_2,*j*_ is the rate of photobleaching of the second population in that cell. We emphasise that these rates are specific to each cell.

### Modelling measurement noise

To infer numbers of molecules, we require a model for the measurement noise generated by fluorescence microscopes.

A common description of this error is additive Gaussian noise [1, 2, 4, 18, 36]. Writing 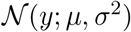 for a Gaussian distribution in *y* with mean *μ* and standard deviation *σ*, *y*_*j*_ (*t*) for the measured fluorescence of cell *j* at time point *t*, and *v* for the fluorescence per molecule - the parameter that we wish to infer, we have 
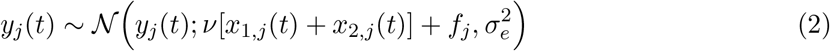
where *j* again indexes an individual cell. Here *σ*_*e*_ is a constant, common to all cells, that determines the magnitude of the measurement noise, and we include a heterogeneous autofluorescence for each cell as *f*_*j*_ (Methods).

The additive Gaussian model for noise has been criticised, and multiplicative Gaussian noise [37] and log-normal noise [3] proposed as alternatives. Unlike additive Gaussian noise, both these models have the intuitively attractive property of a standard deviation that scales with the mean: we do not expect to have the same accuracy when measuring, for example, 100 and 10,000 proteins. This scaling is physically justified if the major sources of measurement noise are inconsistent illumination, segmentation errors, and small movements of the stage. Poisson noise, which is often used to model errors generated by CCD cameras [38], has the similar property that the standard deviation scales with the square root of the mean.

We therefore considered a second model for measurement noise, which we call polynomial state-dependent normal noise, in which the measurement is distributed as: 
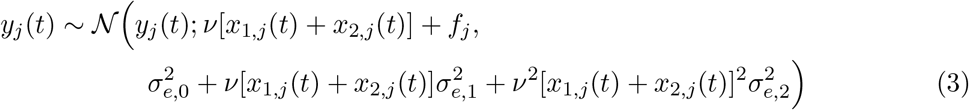
for constant *σ*_*e*,*i*_. This noise model is general: if *σ*_*e*,1_ = *σ*_*e*,2_ = 0, the model recovers Eq. 2; if *σ*_*e*,1_ = 0, the model’s standard deviation scales with the mean. The model does not have the positive skewness of log-normal noise and has support for negative fluorescence values, which can be observed after correcting images for background.

### A fluctuation analysis

Due to their complexity, these models are no longer amenable to previous approaches [18, 21], and we instead use Bayesian methods for the analyses of time-series.

Briefly (see Methods for details), we employ the linear noise approximation [16, 39] to describe fluctuations in the dynamics of bleaching and combine this approximation with a Kalman filter [36, 40] to estimate the likelihood of the parameters. These parameters include homogenous parameters shared between all cells (*v* and *σ*_*e*_) and heterogeneous parameters that vary from cell to cell (λ_1,*j*_, λ_2,*J*_, and *f*_*j*_). To move from this likelihood to inference, we first use an optimisation routine to estimate the parameter values with maximum likelihood and then take these values as a starting point for an adaptive Markov chain Monte Carlo scheme to sample the parameters’ posterior distribution [41].

For additive Gaussian noise (Eq. 2), a standard Kalman filter can be used [40], but for state-dependent noise (Eq. 3), we use the predicted values of the protein concentrations at each time point to find an estimated value for the variance in Eq. 3. As the inference proceeds sequentially, this approximation means that the variance is known at each time point allowing the standard Kalman filter to be employed again, speeding up the analysis.

We verified the algorithm for both types of measurement noise using simulated data and use the evidence for each model, *P*(data|model), to discriminate between models [42].

To compare with biochemical measurements of protein numbers, we use the fluorescence data at the initial time-point before we bleach the cells and the posterior distribution for the parameters of the model given the bleaching data to infer the expected number of proteins in log-phase cells.

### Comparison with biochemical estimates of protein numbers

To test our method, we looked at biochemical measurements of the numbers of molecules for the proteome of budding yeast [15] and selected six proteins (Fus3, Hog1, Guk1, Def1, Gpm1 and Pgk1), which have a range of copy numbers. GFP fusions for these proteins [43] were subjected to our analysis.

The data favour both bi-exponential photobleaching and a state-dependent measurement noise. Running our inference procedure (Methods), we estimated the evidence, *P*(data|model) or the marginal likelihood, for four models: either mono- or bi-exponential bleaching and either additive Gaussian or state-dependent measurement noise. Following standard interpretations of the numerical values of evidence [44], bi-exponential photobleaching is highly favoured and a state-dependent measurement noise preferred over Gaussian noise (Fig. 3A).

**Figure 3.**
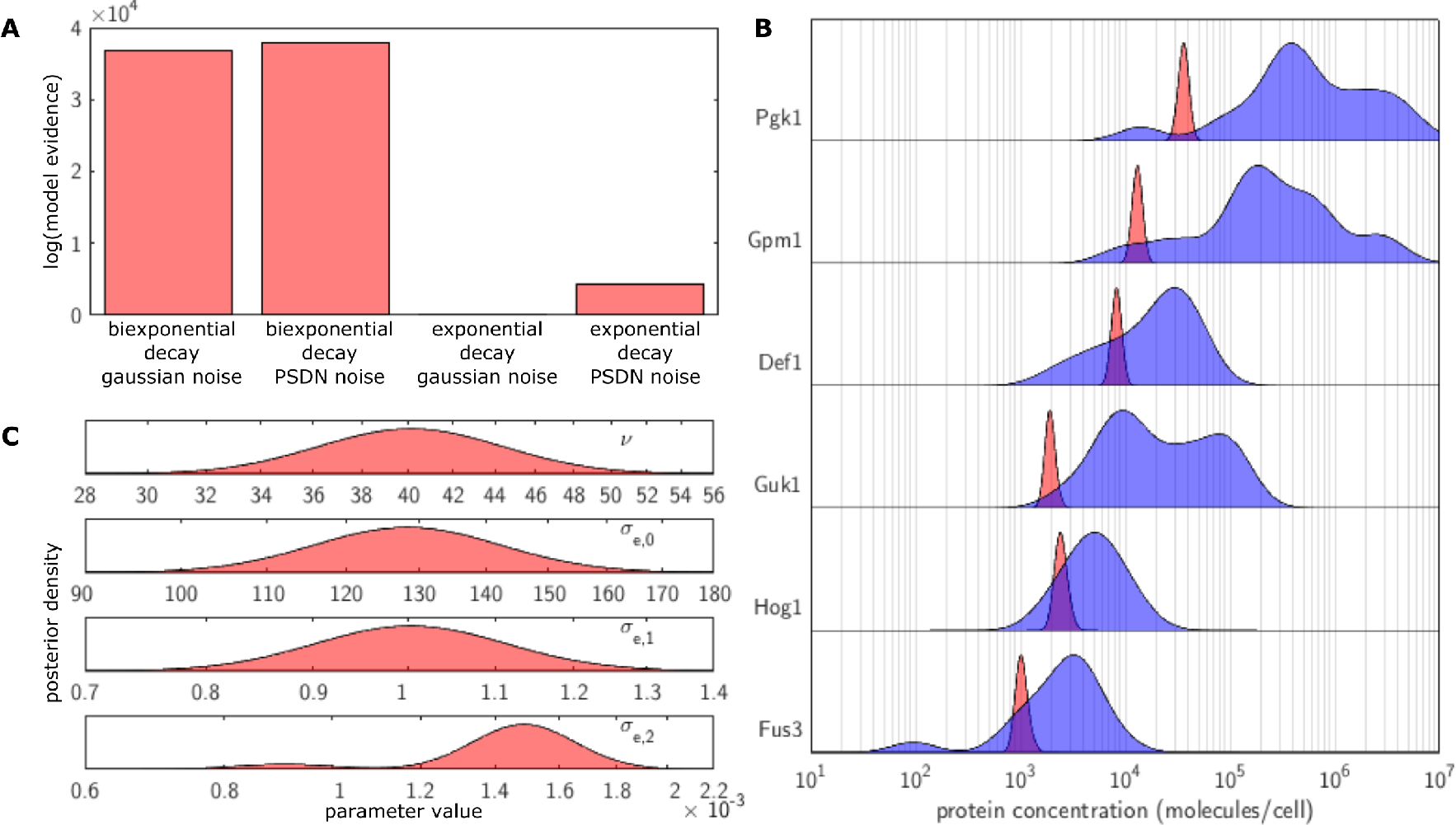
For multiple proteins in budding yeast, estimates of numbers of proteins by our analysis of fluctuations in photobleaching agree with those obtained biochemically. (A) Considering the log evidence for the four models (either mono-or bi-exponential bleaching and either additive Gaussian or state-dependent measurement noise), the data strongly favour bleaching with a biexponential decay and a state-dependent measurement noise (the evidence for this model is at least 5000 dB higher than any other). We normalize so that the lowest log evidence is zero (for the model with mono-exponential bleaching and Gaussian measurement noise). (B) Our analyses, using the favoured model of Eqs. 1 and 3, predicts numbers of proteins in agreement with biochemical estimates. For each protein, the blue histogram is a kernel density estimate of the distribution of protein numbers measured by diverse biochemical methods [15]; the red histogram shows the distribution of inferred numbers for log-phase cells using our analysis (from multiple, replicate photobleaching experiments - Methods). (C) The posterior distributions for the homogenous parameters peak at physically reasonable values and provide, with Eq. 3, a model for the measurement noise of wide-field fluorescence microscopes. The plots show the marginal posteriors (from all replicates - Methods).

Using this model, we infer numbers of proteins in agreement with the biochemical estimates (Fig. 3B). There appears to be a general tendency to underestimate the number of molecules, especially for proteins with high-copy numbers, but there is considerable variation in the biochemical results, which may themselves be biased, and GFP-tagging could affect the stability of a protein and its mRNA.

Understanding the possible reasons the algorithm might give an underestimation requires developing an intuition for how protein number is inferred. A fluorescent measurement, *y*, is given by the product of the brightness per molecule, *v*, and the number of fluorescent molecules, *x*, and is therefore not informative about their individual values. The magnitude of the fluctuations in the underlying biochemical process changing *x* (here photobleaching) are expected to scale with the mean [16]: Var[*x*] ~ *E*[*x*]. Therefore the variance in the fluorescence, *y* = *vx*, scales as Var[*y*] ~ *v*^2^*E*[*x*] = *vE*[*y*]. For a given fluorescence (a given *E*[*y*]), large fluctuations therefore imply a high value of v and correspondingly low numbers of proteins because *E*[*x*] = *E*[*y*]/*v*, and vice versa (Fig. 1). In our algorithm, we make the common assumption that the measurement noise on each measurement is independent. Any correlation in the measurement noise over time will therefore be interpreted as originating from bleaching. The magnitude of the fluctuations in bleaching will correspondingly be overestimated and the number of molecules underestimated. Physically, there are multiple possible sources of correlated error: for example, a gradual drift in the brightness of the illuminating LED or if the microscope slide or cells were to slowly move during the experiment. Furthermore, shortcomings in our biochemical model of bleaching will lead to a discrepancy between the model’s expected behaviour and the cells’ actual behaviour that would appear too as correlated errors.

Our method not only returns an estimate for the number of molecules but also posterior probabilities for the parameters necessary to describe a wide-field fluorescence microscope’s distribution of measurement noise (Fig. 3C). It is difficult to find published data with which to compare these parameters. Nevertheless, the range of supported *σ*_*e*,1_ values is close to 1, which is the value expected for Poissonian noise. These estimates are robust: using an informative prior for the brightness parameter, *v*, the modes of the posterior distributions maintain the same order of magnitude and are shifted at most by a factor of two (Methods).

## Discussion

A long standing problem in quantitative fluorescence microscopy is converting measurements of fluorescence to absolute units. Absolute units enable the data from different laboratories to be combined and are needed both for models to accurately fit single-cell data [3] and to assess the validity of the magnitude of fitted parameters [45]. As such, absolute units are an important facilitator for the long-term success of systems and synthetic biology [46, 47].

Here we have presented a fluctuation analysis that estimates the numbers of fluorescent proteins by deliberately photobleaching cells on a wide-field fluorescence microscope. Fluctuation analyses have long been used to estimate numbers of molecules on confocal microscopes, but the challenges of untangling measurement noise from fluctuations in stochastic biochemistry has prohibited a similar approach becoming a standard for the more commonly used wide-field fluorescence microscope.

Building on advances in the study of stochastic gene expression [40], we have used the linear noise approximation to model stochastic biochemical events combined with a Kalman filter and a state-dependent measurement noise to infer numbers of proteins from fluctuations in photobleaching. For budding yeast, we have shown that photobleaching has two distinct time-scales, corresponding to two decay processes, but our framework remains valid for more complex reactions. The approximations developed allow faster inference than alternative approaches, such as particle-based methods, and the data we require can be gathered on a standard wide-field fluorescence microscope, requiring no specialist equipment.

Further, we have rigorously compared different models of measurement noise, and we anticipate that the state-dependent measurement noise, Eq. 3, which is most consistent with our data, together with its inferred parameters (Fig. 3C), will underpin future work both for fitting mathematical models and for quantitative microscopy [48].

To bring together discoveries from different laboratories and to build those discoveries into a larger predictive framework, measurements both in single cells and in absolute numbers are necessary. We believe that methods that extend the fluctuation analyses available for confocal microscopes to wide-field microscopes, such as the one we present here, are key for a broad transition to a quantitative single-cell biology.

## Methods

### Selecting proteins to study

When selecting proteins to test our method, we looked at three whole-proteome datasets of absolute numbers of protein obtained by quantitative Western blot [26], mass spectrometry [14], and fluorescence microscopy [49]. Proteins were selected that appeared in all three data sets. To obtain proteins whose levels would be robust to any stresses from our fixing procedure, we used a protein localisation atlas [50] to select cytoplasmic proteins that showed no significant fold change under starvation and dithiothreitol-and peroxide-induced stress. From these proteins, we selected four giving a broad range of levels: Def1, Guk1, Gpm1, and Pgk1. To these proteins we added Hog1 because of its regular study in our laboratory [51] and Fus3 because of the availability of a measurement by fluorescence correlation spectroscopy (FCS) [52]. Taking a cytoplasmic concentration of 180 nM for vegetative *Saccharomyces cerevisiae* cells and a three times higher concentration for the nucleus [52] along with cellular and nuclear volumes of 42 *μl* and 3 *μl* [53, 54, 55], we estimate 5200 molecules of Fus3 per cell from this FCS data.

During the course of our work, however, a comprehensive collection of whole-proteome data sets was published [15], and this data is the data we use for comparison. Our estimate for Fus3 from the FCS data is within the range of values reported in this data set, being 1.75 times the median.

### Cell preparation

Following [26], cells from the ORF GFP collection [43] were grown overnight in YEPD (2%) media past the diauxic lag, and 0.5 ml of this culture diluted in 5 ml of fresh media. After 5 hours and at an OD of ~ 0.5, the cells were fixed [56].

### Microscopy and image analysis

All experiments were performed on a Nikon Eclipse Ti inverted microscope controlled using custom MATLAB scripts (Mathworks) written for Micromanager [57] and the Perfect Focus System. We used a 60X 1.2 NA water immersion objective (Nikon), and images were acquired using an Evolve EMCCD camera (Photometrics) with a 512 × 512 sensor in CCD mode. Cells were adhered to slides using concanavalin A.

To effect photobleaching, the GFP excitation LED was kept at full power for the duration of the experiment with cells imaged for GFP every 10 seconds. This procedure was repeated for multiple, isolated fields of view for each slide. Wild-type cells, which do not express GFP, were also imaged.

Fluorescence images were corrected for flat field and background. We obtained a flat field image by flowing 0.001% fluorescein (by mass) through a narrow microfluidic device [58] and imaging multiple positions over an extended time course. Any microfluidic features were ‘blotted out’ of the images and the modified images averaged and normalised to a median of one. A correction was applied to the fluorescence images through an element-wise division by the flat field. To remove any background particular to the slide, we subtracted the average pixel fluorescence for a region of the image containing no cells from each pixel value (Fig. 4A). Fluorescence images were also registered to correct for drift in the field of view.

**Figure 4.**
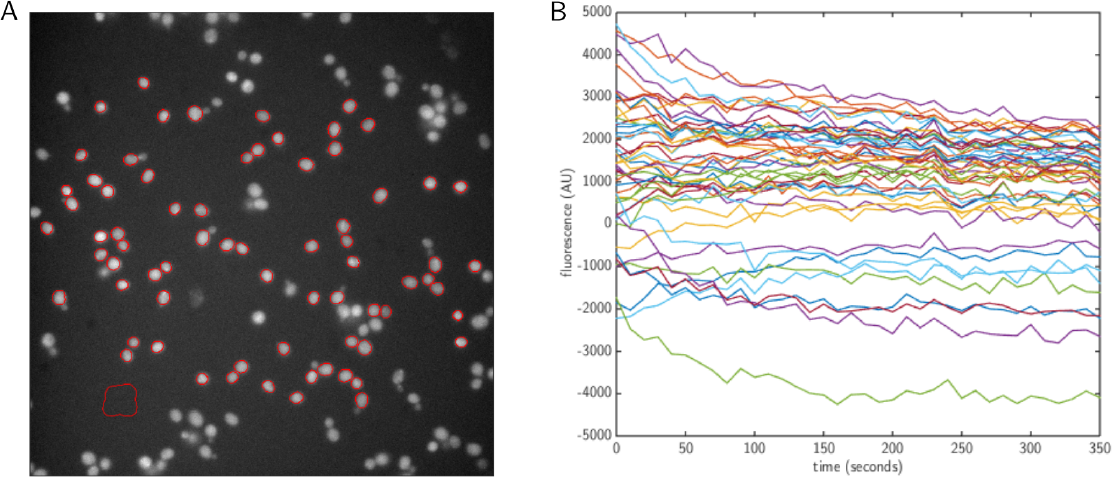
After correction for flat field and background, cells have a residual autofluorescence. (A) Examples of selected cells with their outlines in red. The area in the bottom left without cells was used to correct the fluorescence for background. (B) The residual fluorescence of each wild-type cell, after all corrections, has a non-zero mean, which we include in our models of measurement noise, Eqs. 2 and 3, as the heterogenous parameter *f*_*j*_.

Given that we imaged fixed cells, cells were selected and segmented at the first time point from a bright-field image and whole-image registration used to propagate this outline to other time points. Cells in the bright-field image were chosen by eye to be isolated, well focused, present for the whole experiment (i.e. not washed away), and in a region where the illumination intensity (taken from the flat field correction) was at least 80% of the median illumination. Selected cells were outlined based on the out-of-focus bright-field image using custom MATLAB scripts, and these outlines curated by hand.

Cells were also corrected for autofluorescence. We corrected wild-type cells for flat field and background, and their average value was subtracted from all fluorescent cells. When this procedure is performed on wild-type cells, the residual fluorescence time-series display a systematic error that varies from cell to cell (Fig. 4B). We therefore assume a similar systematic error in the time-series of our fluorescent cells and include a heterogeneous basal fluorescence in our model of measurement noise (*f*_*j*_ in Eqs. 2 and 3).

Finally, cell fluorescence was calculated as the sum of the values of brightest 80% of pixels within the cell boundary. This measure reduced the effect of movements of the stage. We used the 35 time points after the 5th because the first 5 time points were often erratic. Cells or replicates that displayed large systematic deviations from bi-exponential decay (such as a sudden drop in fluorescence) were discarded.

We twice repeated the entire experimental procedure (bringing colonies up from glycerol stocks, cell fixation and imaging). The list of all replicates used, with cell numbers per replicate, are shown in Table 1.

**Table 1.** Numbers of cells for each experimental replicate for each protein. On two occasions, cells were brought up from stocks, cultured, fixed, and photobleached. Cells were imaged for 400 ms except for asterisked replicates, which were imaged for 200 ms to avoid saturating the camera. We compensate for this difference when combining estimates (Methods).

| Date | Strain | $n$ for each replicate |
| --- | --- | --- |
| 11 Oct 2016 | Def1-GFP | 30, 30, 30 |
|  | Fus3-GFP | 30, 25, 30 |
|  | Gpm1-GFP | 17, 30 |
|  | Guk1-GFP | 30, 30, 30 |
|  | Hog1-GFP | 8, 17, 30 |
|  | Pgk1-GFP | 9*, 21*, 12* |
| 23 Mar 2017 | Def1-GFP | 30, 30 |
|  | Fus3-GFP | 30, 30, 30, 30, 30 |
|  | Gpm1-GFP | 30, 30, 30, 30 |
|  | Guk1-GFP | 30, 30 |
|  | Hog1-GFP | 24, 24, 30, 30 |
|  | Pgk1-GFP | 30, 30, 30, 30 |

### Inference fails assuming no measurement noise and mono-exponential decay

Analysing the data using the previously proposed estimator [21], which assumes no measurement noise and that photobleaching decays exponentially, the number of proteins inferred are inconsistent with the biochemical data. To correct for residual autofluorescence, we first fitted an exponential decay with an offset to each cell and then subtracting the fitted offset from the data before processing (a procedure that worked for data simulated from a single exponential decay with parameters shared by all cells and no measurement noise). Other than this correction, we followed the procedure of Kim *et al.* [25].

This naive inference underestimated numbers of proteins by several orders of magnitude. For Fus3, 174 and 1 proteins are estimated for the two replicates (the biochemical estimate is 3835 [15] - an underestimate by at least a factor of 20); for Hog1, the estimates are 69 and 77 (an underestimate by a factor of 90); for Guk1, the estimates are 153 and 1 (an underestimate by a factor of over 420); for Def1, the estimates are 7 and 47 (an underestimate by a factor of over 750); for Gpm1, the estimates are 134 and 11 (an underestimate by a factor of over 4500); for Pgk1, the estimates are 34 and 9 (an underestimate of over 45000).

### Inference with measurement noise and complex decay

Given data **Y**_*t*_ = {**y**_1_, ⋯, **y**_*t*_}, we would like to infer ***θ*** = {***θ***_*m*_, ***θ***_*e*_}, where ***θ***_*m*_ are the parameters for the biophysical model and describe the dynamics of the underlying **x** variables and ***θ***_*e*_ are the parameters describing the distribution of the measurement noise (including the autofluorescence *f*_*j*_). The **y** variables are related to the **x** variables through this measurement noise. We note that an analytical, maximum likelihood solution is possible for the ideal case without measurement noise [56].

### Modelling protein concentration using the linear noise approximation

The dynamics of the protein concentrations, **x**, are determined by chemical reactions and can be described by a chemical master equation for *P*(**x**, *t*), the probability distribution of the state **x** at time *t* [16]. We use the linear noise approximation (first-order terms in an expansion of the master equation in the size of the system, which for us is the volume of the cell [39]). This approximation makes *P*(**x**, *t*) a normal distribution if the initial distribution is either a normal or a delta distribution [16].

If we let the stochiometric matrix be **S** and the hazards (propensities) be **h** and noting that **x** describes numbers of molecules not concentrations, then 
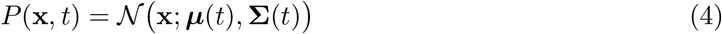
where ***μ***(*t*), the mean, and ∑(*t*), the covariance matrix, obey [40] 
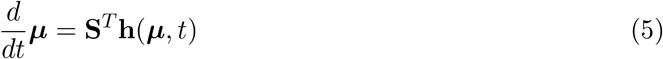
and 
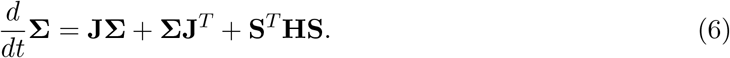

Here **J** is the Jacobian: 
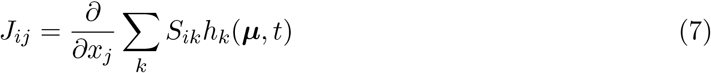
and **H** is a matrix of zeros with **h**(***μ***, *t*) on the diagonal.

### Linear noise and sequential data – a Kalman filter

We wish to infer ***θ*** given **Y**_*t*_. Bayes’s rule states: 
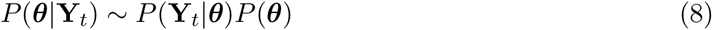
or, by using the rules of probability to factorize the likelihood, 
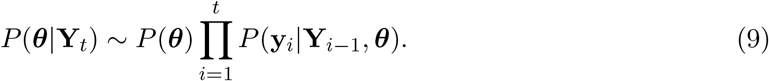

We find each term in Eq. 9 sequentially by considering the dynamics of **x** from one time point to the next and then correcting that dynamics given the observed data [40]. Assume that at time point *i* − 1 the distribution *P*(**x**_*i*−1_|**Y**_*i*−1_, ***θ***) is Gaussian with a known mean 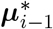 and covariance matrix 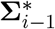: 
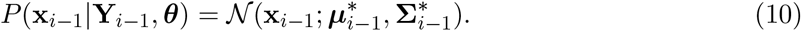

Using the linear noise approximation to describe the dynamics of **x**, we can, with Eq. 10 providing the initial condition, integrate Eqs. 5 and 6 over one time interval to time point *i* to find ***μ***_*i*_ and ∑_*i*_ and that 
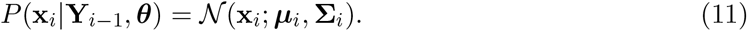

We next wish to extend the conditioning in Eq. 11 to include the data point, **y**_*i*_, at time point *i*. To do so, note that 
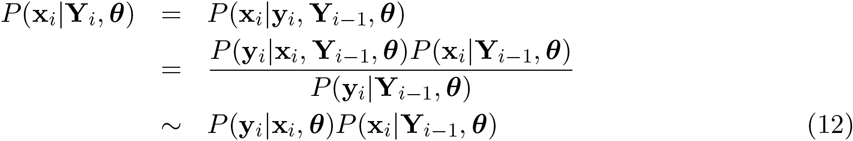
using Bayes’s rule, conditioning on **Y**_*i*−1_ and ***θ***, and assuming that the measurement noise only depends on the current value of **x**. If the measurement noise too has a normal distribution 
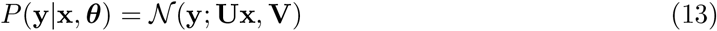
with **U** being a constant projection matrix and **V** being a covariance matrix, then we can simplify Eq. 12 using the properties of normal distributions [59]. We find that *P*(**x**_*i*_|**Y**_*i*_, ***θ***) is also normal with a mean 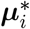 and a covariance 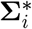 that satisfy [40] 
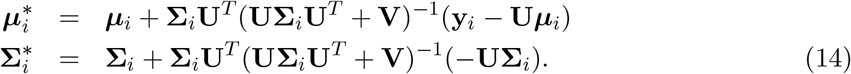

We see that the prediction of ***μ***_*i*_ and ∑_*i*_ found from 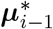 and 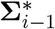 using the linear noise approximation are corrected because of the new data point and the measurement noise.

The factors in Eq. 9, *P*(**y**_i_|**Y**_*i*−1_, ***θ***) obey 
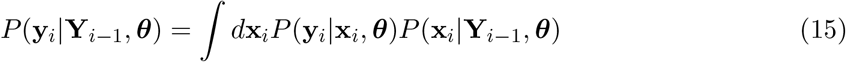
and are therefore the normalizing factors for Eq. 12, satisfying 
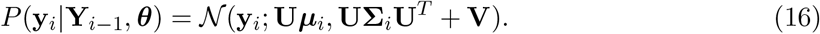

Hence from a normal prior distribution for **x**_1_, *P*(**x**_1_|***θ***), we use Eq. 15 to find *P*(y_1_|***θ***), the first term in the factorization of the likelihood in Eq. 9, and Bayes’s rule to find *P*(**x**_1_|**y**_1_,***θ***) ~ *P*(**y**_1_|**x**_1_,***θ***)*P*(**x**_1_|***θ***), the starting normal distribution in Eq. 10 for the sequential inference.

To specialize the algorithm to photobleaching, we subtract the autouorescence, *f*_*j*_, from each data point before applying the Kalman filter. The Kalman update step, Eq. 14, can result in unphysical, negative components of ***μ***, which we set to zero and then continue the Kalman filter.

### Application to biophysical models of photobleaching

To explain the observed biexponential decay during bleaching, we consider two populations of molecules that bleach at different rates 
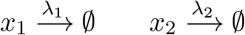
for which we can analytically solve Eqs. 5 and 6.

For this model 
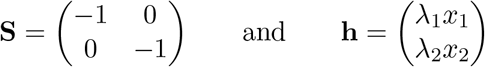
and 
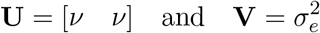
in Eq. 13.

Eq. 5 becomes 
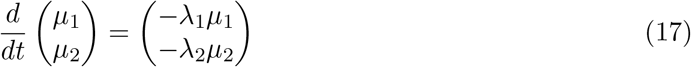
so that 
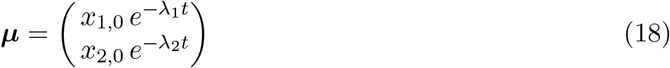
for initial condition **x**_0_. In Eq. 6, 
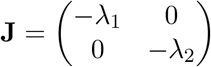
and 
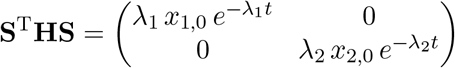
which upon integration implies 
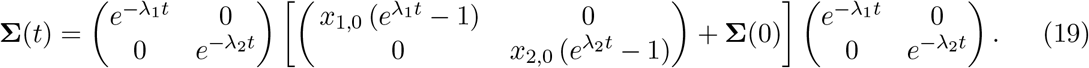

### Starting the inference

To begin the inference scheme, we must first specify a prior *P*(**x**_1_|***θ***_*m*_). To do so, we introduce two new parameters: *x*_0_, which is the total amount of fluorescent protein at time point 0, and *α*, which is the partitioning of this fluorescent protein between the slow-and fast-bleaching pools. We infer both these parameters.

For the Kalman filter, we require that *P*(**x**_1_|***θ***_*m*_) be a normal distribution, but we wish to start **x**_1_ at a known value. We therefore assume that 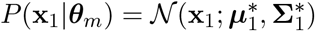, with 
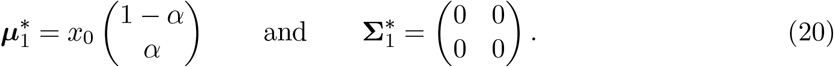

This distribution is a delta function because 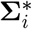 is a matrix of zeros and is not strictly normal. Nevertheless, Eq. 20 imposes the known initial value 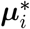.

### Sampling the posterior probability

To sample from the posterior probability, *P*(***θ***|**Y**_*t*_), in Eq. 8 we use both optimization and a Markov chain Monte Carlo method.

We distinguish between heterogenous parameters, which are specific to each cell (λ_1_, λ_2_, *f*, *x*_0_ and *α*), and homogeneous parameters, which have the same values for all cells (*σ*_*e*_ and *v*). The homogenous parameters and *x*_0_ have scale-free priors: for example, *P*(*v*) = 1/*v*. The heterogenous parameters other than *x*_0_ have flat priors. All priors are proper and bounded to reasonable physical values. To ensure the expected behaviour is dependent only on the heterogeneous parameters and to improve the mixing of the Markov chain, we propose the combination *vx*_0_, referred to as *b*_0_, rather than *x*_0_.

To generate samples from the posterior, we use a Metropolis-within-Gibbs scheme [60] with the heterogeneous parameters updated separately from the homogeneous parameters. We employ adaptive parallel tempering to accelerate mixing in the Gibbs sampler [61], which has been found to perform well on benchmark biochemical models [62]. Specifically, we use 10 chains with their temperatures chosen adaptively. Parameters are proposed independently, with λ_1_, λ_2_, *f* and *α* proposed from a normal distribution and *b*_0_, *σ*_*e*_ and *v* proposed from a log-normal distribution. The scales for the proposal distributions for the heterogeneous and homogeneous parameters are adaptively selected.

To start the Monte Carlo method, we try to find values of the parameters that maximize the likelihood *P*(**Y**_*t*_|***θ***). We use a nested optimisation scheme [37]:

1. We find starting values for the heterogeneous parameters by fitting a bi-exponential decay to each cell’s time series.
2. All parameters, including the heterogenous parameters, are then fitted for each cell, independently of all other cells, using a particle swarm.
3. We create an initial parameter set for the homogenous parameters by taking the median of the homogeneous parameters from the individual fits of Step 2.
4. We perform iterative optimisation: first optimising the homogeneous parameters with the heterogeneous parameters fixed then vice versa until a maximum is reached. Each individual optimisation used Matlab’s fmincon with the default algorithm.
5. To provide diverse starting points for the chains, half of our chains are initialised at the parameter values found in Step 4 and the other half are initialised from optima found by performing the iterative optimisation of Step 4 from random parameters rather than from those found in Step 3.

### Extending to state-dependent measurement noise

The variance of the measurement noise of Eq. 3 depends on the state of the system — the **x**_1,*j*_ and **x**_2,*j*_. Our inference scheme, however, assumes a constant variance and so is no longer valid.

We therefore approximate Eq. 3 by replacing the explicit state dependence in the variance by its expected value given **Y**_*t*−1_: 
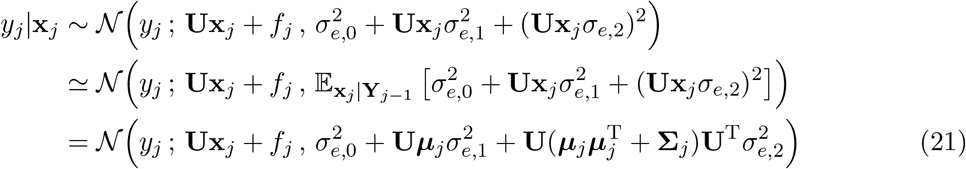

The usual update step in the Kalman filter, Eq. 14, can then be used.

This estimate of the likelihood performs comparably with a particle filter and is more robust [56]. We use the Metropolis-within-Gibbs scheme to sample the posterior probability with lognormal proposal distributions and scale free priors for the additional homogeneous parameters: *σ*_*e*,1_ and *σ*_*e*,2_.

#### *In silico* verification

We verified our methodology using simulated data (Fig. 5). For additive Gaussian measurement noise, the inference is able to accurately infer *v* if the measurement noise is not so high as to mask the stochastic fluctuations. This threshold depends on the initial number of proteins. For any magnitude of measurement noise, there will be a sufficiently high initial number that stochastic fluctuations are detectable. For state-dependent noise, however, the inference no longer necessarily improves with increasing numbers of protein when the state-dependent noise is sufficiently large (data not shown).

**Figure 5.**
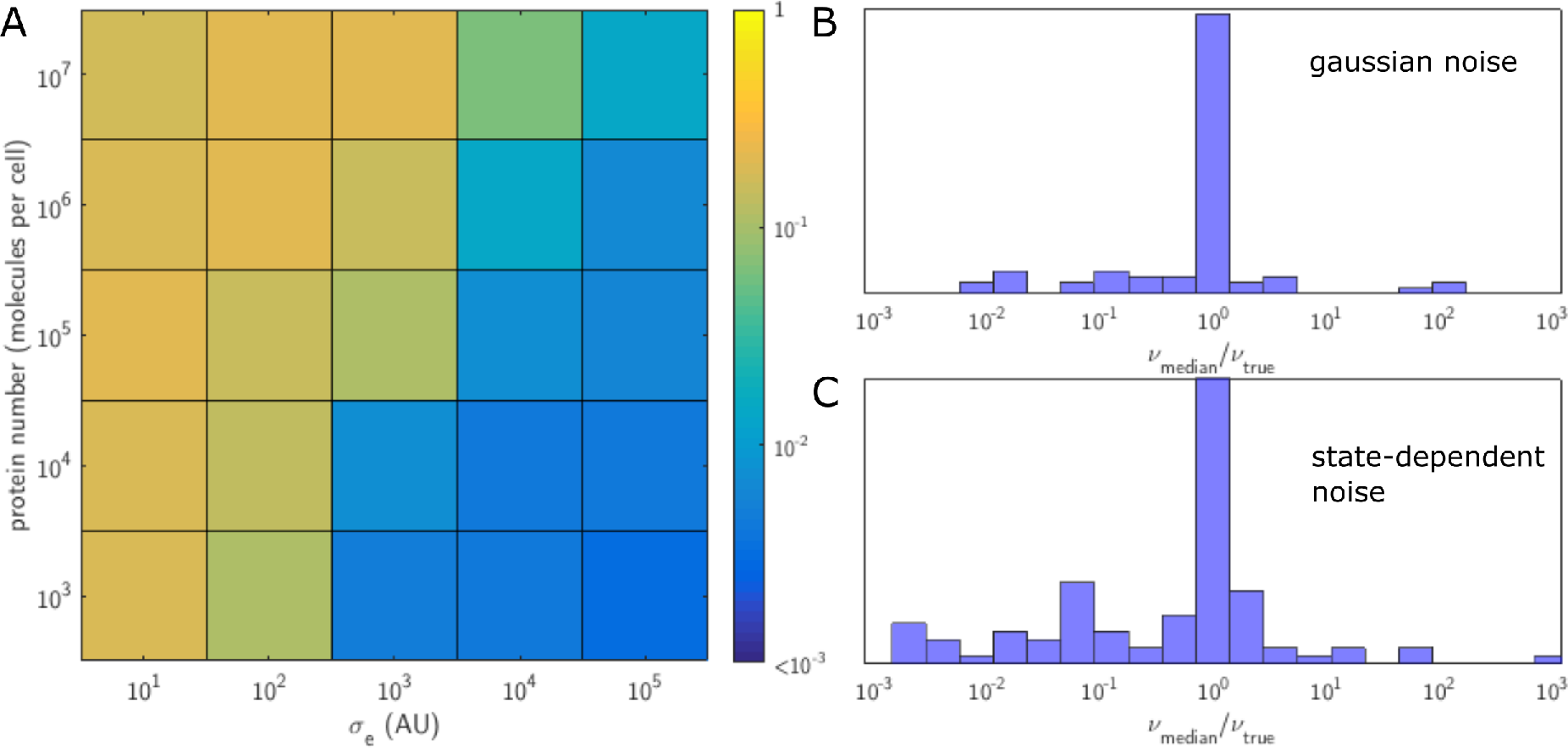
Parameters are well identified for simulated data. We simulated time-series for the model of Eq. 1 with a distribution of decay rates and initial numbers of proteins similar to those observed experimentally. (A) For Gaussian measurement noise, we show the probability of the true value of *v* averaged over 5 simulated datasets (and normalized to correct for the kernel density estimates we use - Methods). A value of 1 would indicate total confidence in the true value. When the initial number of proteins (*y*-axis) is large compared to the measurement noise (*x*-axis), *v* can be accurately inferred. Here *v* = 10. (B) Letting *v* = 1 and *v* = 100 too, we show the results for all three values of *v* as a histogram of log_10_ (*v*_median_/*v*_true_). The spike at 1 indicates v can usually be correctly inferred. (C) A similar histogram for state-dependent measurement noise also shows that inference of *v* typically remains accurate if more challenging.

### Estimating the expected numbers of proteins

Given a sample of parameters from the posterior distribution *P*(***θ***|**Y**), and a fluorescence measurement *y*, we would like to find the posterior distribution of the number of proteins, *x*: 
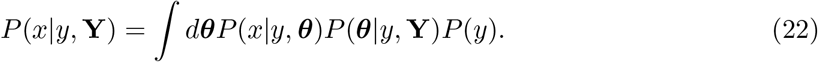

We assume that the measurement *y* does not change the posterior probability of ***θ***: *P*(***θ***|*y*, **Y**) ≃ *P*(***θ***|**Y**). Ignoring *P*(*y*), which is independent of *x*, we have 
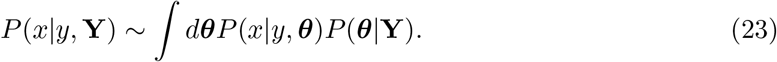

By Bayes’s rule 
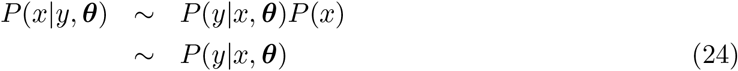
assuming a constant prior, *P*(*x*).

For Gaussian measurement noise, Eq. 2, we use that 
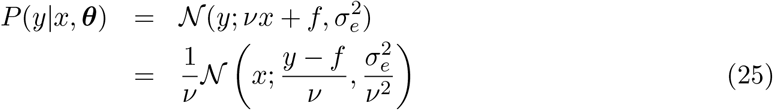
from the properties of Gaussian distributions. From Eq. 24 then 
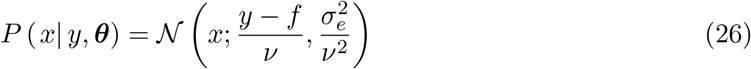
normalizing over *x*.

For state-dependent measurement noise, Eq. 3, we have 
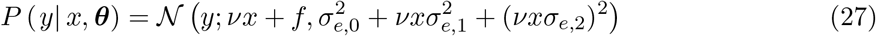
which cannot be straightforwardly inverted to give a normalized distribution in *x*. We therefore approximate *vx* by *y* so that the variance in Eq. 27 becomes constant. Then 
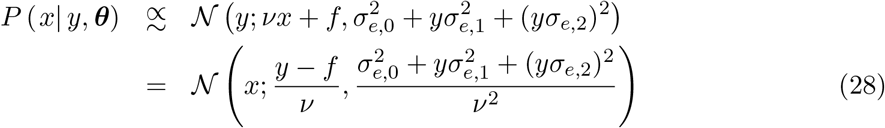
similarly to Eq. 26.

Continuing with 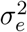 as the variance for simplicity (although the results will hold too for the constant variance in Eq. 28), Eq. 23 becomes 
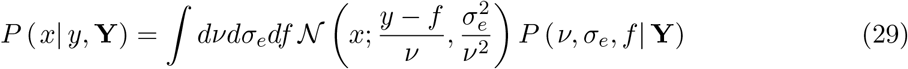
which we estimate on a grid of *x* values as an average over the Monte Carlo samples generated from *P* (*ν*, *σ*_e_, *f*| **Y**).

#### Multiple measurements

If we have multiple, replicate measurements of fluorescence (each, for example, from a different cell) and are interested in the distribution of *x̄* = ∑*x*_*i*_/*N*, we can use that the distribution of a sum of Gaussian variables is also Gaussian with a mean and a variance equal to the sum of the mean and variances of all the Gaussian variables in the sum [16]: 
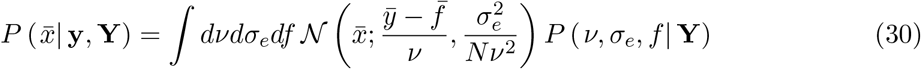
following Eq. 29. Practically, we assume that 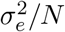 and *f̄*/*ȳ* sufficiently small that we can take the limiting case: 
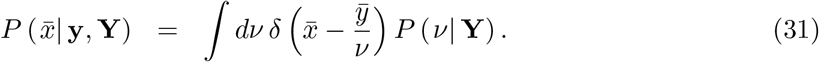

We use a kernel density representation for *P* (*x̄*| y, **Y**) with kernels 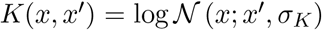 and 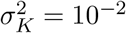: 
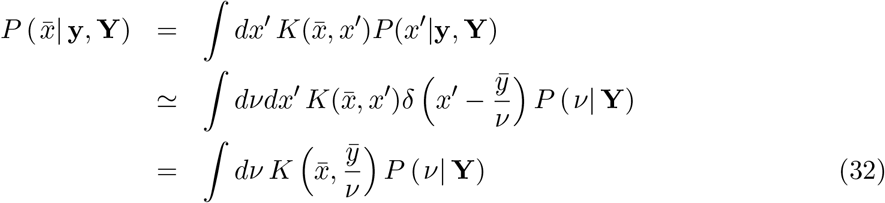
using Eq. 31. We evaluate the integral in Eq. 32 using the Monte Carlo samples of *P* (*v*| **Y**).

### Combining estimates from all datasets

To have a final estimate of the homogeneous parameters for analysing other microscopy experiments, we combined posteriors from all replicates for all genes (the posteriors in Fig. 6). Writing 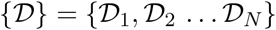 as the set of data from all replicates, we wish to sample from 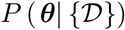. We note that the datasets are conditionally independent given ***θ*** so that 
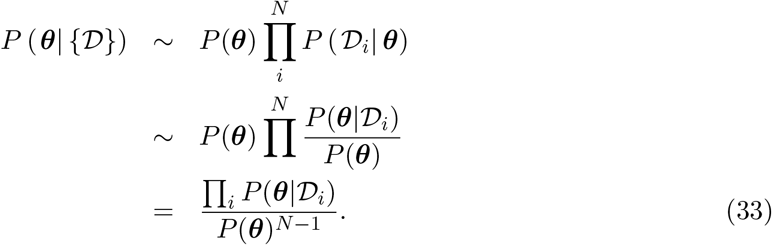

**Figure 6.**
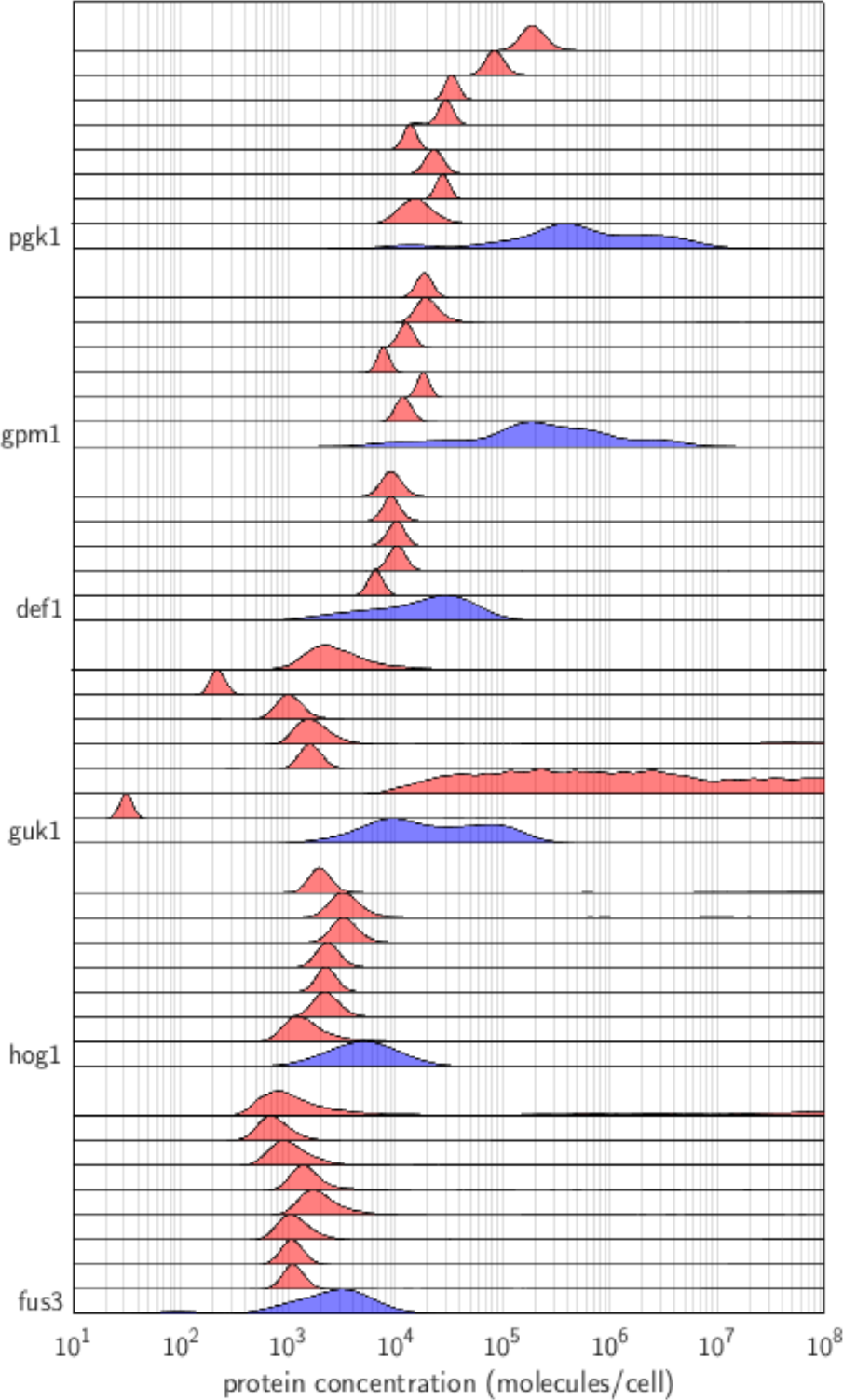
Inference from individual replicates is typically consistent. For each protein, the blue histogram is a kernel density estimate of the distribution of protein numbers using different biochemical measurements [15]; the red histograms show the distribution of inferred numbers using our analysis for each replicate.

We approximate the 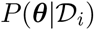 in Eq. 33 with kernel density estimates. If {***θ***_*i*,1_,… ***θ***_*i,M*_} is the set of parameter samples from the posterior 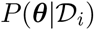 obtained from our Monte Carlo method, then 
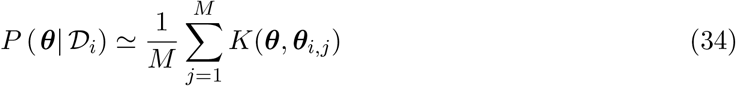
where the kernel functions are isotropic log normal distributions with variance 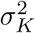 as before. In principle, these kernel density estimates allow Eq. 33 to be evaluated at any ***θ***; in practice, we use a Metropolis algorithm to sample ***θ*** to overcome the dimensionality of the parameter space.

This procedure was used to both combine the posteriors for replicate datasets (Fig. 6) for a given protein (Fig. 3B) and combine the posteriors for all replicates and all proteins (Figs. 3C & 7).

**Figure 7.**
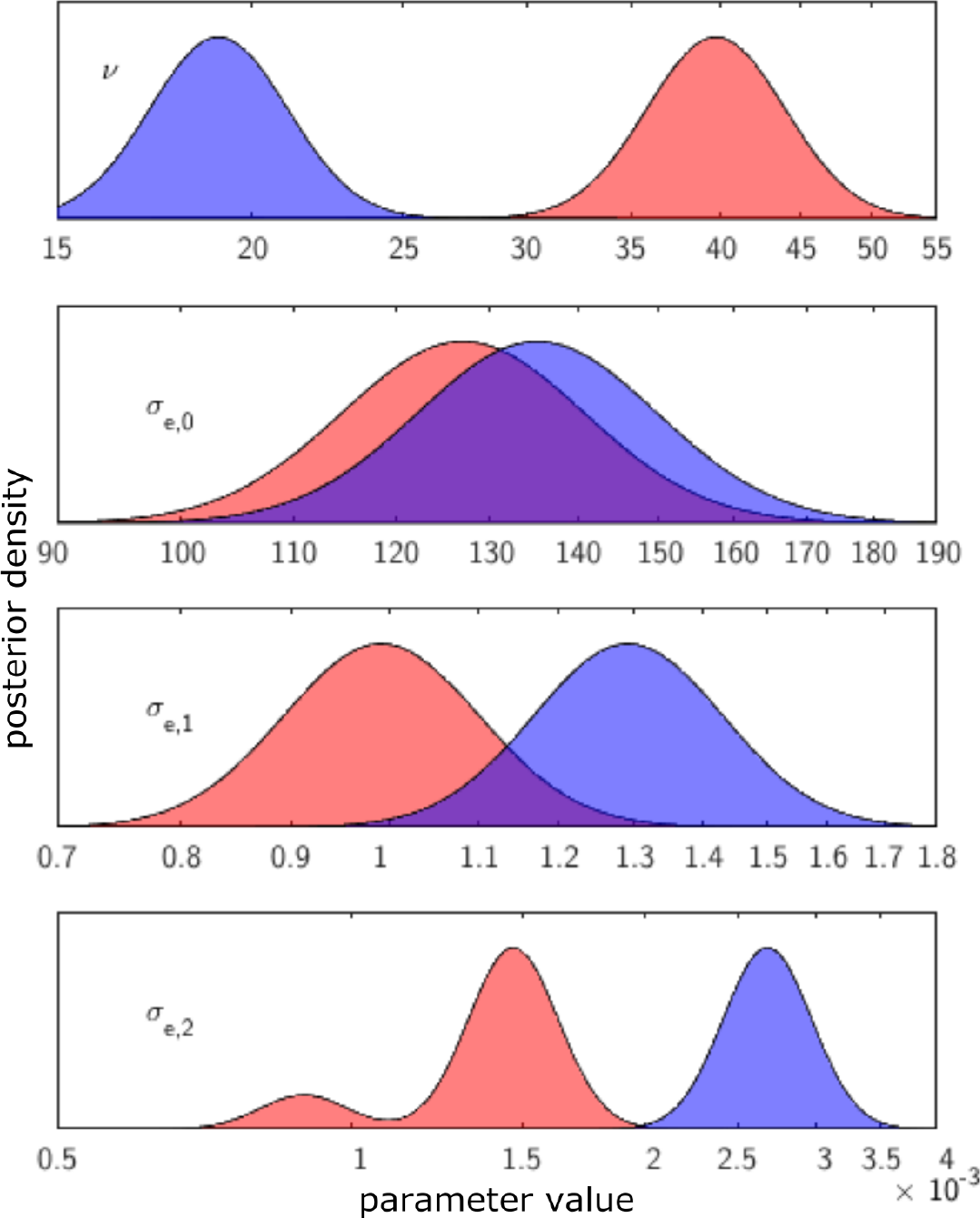
Tightly restricting the *a priori* range of v does not substantially change the inference. The locations of the modes of the posterior distributions for the data of Fig. 3 maintain the same order of magnitude even when this data is used to impose an informative prior for the fluorescence per molecule (distributions with the informative prior in blue; distributions from Fig. 3C in red).

For Pgk1-GFP, we accounted for the difference of a factor of two in the exposure time used in the two biological replicates (Table 1). Before combining samples, the samples of *v* were doubled for the three replicates obtained with the low exposure time. Similarly, we doubled the cellular fluorescence for these three replicates before using the full posterior for *v* to estimate protein numbers.

### Estimating the evidence for each model

To compare models of the measurement noise, we calculate Bayes factors, i.e. a ratio of the log evidence, log *P*(data|model), for four models — either mono-or biexponential bleaching and either Gaussian or state-dependent measurement noise. For a given model, 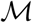, we estimate the evidence, 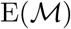, as follows:

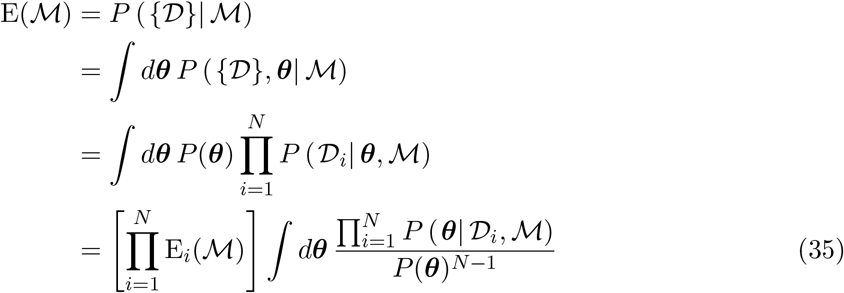
where 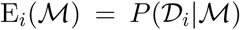 is the model evidence for the data set 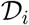. We approximate the individual evidences, 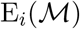, using the Monte Carlo samples for the individual datasets. The second term we approximate using our samples for combining posteriors and the kernel density estimate (Eqs. 33 and 34). In both cases, we use the Laplace approximation to find the evidence and the covariance of the samples to estimate Hessians [42].

In addition to being favoured by the total evidence for all datasets, the bi-exponential bleaching model with state-dependent noise has the highest evidence in the majority of data sets: 25 compared to 8 for bi-exponential decay and Gaussian measurement noise, 6 for mono-exponential decay and Gaussian noise, and 2 for mono-exponential decay and state-dependent noise.

### An informative prior results in only a minor change to the inference

To further test the validity of our model, we repeated the analysis, imposing an informative log-normal prior for the parameter *v*. In so doing, we integrate all available data to obtain a best estimate of all the parameters in our model. These estimates can be used in analysing future experiments, for simulations of microscopy data, and can be compared with the estimates obtained with uninformative priors to assess the reliability of the model. Further, we can again calculate the evidence for each model to identify, given all the information available, the best model for our data.

We used an informative log-normal prior for the parameter *v*, which is the fluorescence per molecule, with parameters fitted to the published data of Ho *et al.* [15]. For each data set, this parameter *v* was confined to be within 1 interquartile range of the median published value: the mode of the prior was set to the median of the published results and the scale factor set to half of their interquartile range.

With the informative prior, the bi-exponential model with state-dependent noise is still selected with a significant Bayes factor (> 10^4^ dB compared to the next best model). The modes of the posterior distributions for the parameters are shifted at most by a factor of 2 (Fig. 7), which is negligible given that the correct order of magnitude is typically all that is required. As expected, the parameter v is shifted down to amend the underestimation of protein numbers, and the measurement noise parameters (*σ*_*e*,2_ and to a lesser extent *σ*_*e*,1_) are increased to accommodate the fluctuations in fluorescence that are no longer explained by the now reduced stochastic fluctuations.

### Data availability

Data generated in this work is available at http://dx.doi.org/.

## Acknowledgements

We would like to thank Ivan Clark, Iain Murray, Ted Perkins, Julian Pietsch, Christos Josephides, and particularly David Schnoerr for comments and suggestions.

## Author contributions

EB performed the experiments and developed the analysis; EB & PSS wrote the paper.

